# Genome annotation with long RNA reads reveals new patterns of gene expression in an ant brain

**DOI:** 10.1101/2021.04.20.440671

**Authors:** Emily J. Shields, Masato Sorida, Lihong Sheng, Bogdan Sieriebriennikov, Long Ding, Roberto Bonasio

## Abstract

Functional genomic analyses rely on high-quality genome assemblies and annotations. Highly contiguous genome assemblies have become available for a variety of species, but accurate and complete annotation of gene models, inclusive of alternative splice isoforms and transcription start and termination sites remains difficult with traditional approaches. Here, we utilized full-length isoform sequencing (Iso-Seq), a long-read RNA sequencing technology, to obtain a comprehensive annotation of the transcriptome of the ant *Harpegnathos saltator*. The improved genome annotations include additional splice isoforms and extended 3’ untranslated regions for more than 4,000 genes. Reanalysis of RNA-seq experiments using these annotations revealed several genes with caste-specific differential expression and tissue-or caste-specific splicing patterns that were missed in previous analyses. The extended 3’ untranslated regions afforded great improvements in the analysis of existing single-cell RNA-seq data, resulting in the recovery of the transcriptomes of 18% more cells. The deeper single-cell transcriptomes obtained with these new annotations allowed us to identify additional markers for several cell types in the ant brain, as well as genes differentially expressed across castes in specific cell types. Our results demonstrate that Iso-Seq is an efficient and effective approach to improve genome annotations and maximize the amount of information that can be obtained from existing and future genomic datasets in *Harpegnathos* and other organisms.

## Introduction

Improved sequencing technologies have enabled studies in previously inaccessible organisms, but annotations remain the bottleneck to thorough genomic and epigenomic analyses. Specifically, gene annotations of many non-model organisms suffer from the limitations imposed by their reliance on traditional, short-read RNA-seq coupled with gene prediction software (Haas et al., 2013; Haas and Zody, 2010; Martin and Wang, 2011). This approach can identify splice junctions but cannot capture complex combinations of exons that define full transcript isoforms. Furthermore, local fluctuations of RNA-sequencing coverage can make it difficult to identify the 5’ and 3’ untranslated regions (UTRs), resulting in inaccurate transcription start sites (TSS) and transcription termination sites (TTS) (Steijger et al., 2013). Inaccurate annotation of the 3’ end of genes is especially problematic for the analysis of droplet-based single-cell RNA-sequencing data, such as those obtained with the widely-used 10X Genomics 3’ gene expression platform (Zheng et al., 2017) or Drop-seq (Macosko et al., 2015), both of which have a strong 3’ bias due to the capture of transcripts using an oligo-dT sequence (Zhang et al., 2019).

Just as long-read sequencing of DNA has led to better genome assemblies (Koren et al., 2017; Shields et al., 2018), long-read sequencing of RNA molecules can be used to address the limitations of short-read-based annotation. PacBio Single Molecule Real-Time (SMRT) sequencing for RNA, called Iso-Seq, produces full-length transcript sequences, resolving issues of isoform reconstruction and accurate identification of the end of transcripts. Iso-Seq has been used in many settings and organisms to reveal alternative polyadenylation sites (Gordon et al., 2015), provide insight into alternative splicing (Feng et al., 2019), and identify tissue-specific transcriptional isoforms (Wang et al., 2016). The genome of the ant *Harpegnathos saltator* was first sequenced in 2010 (Bonasio et al., 2010), was improved using Pacific Biosciences (PacBio) long-read DNA sequencing in 2018 (Shields et al., 2018), and re-annotated by NCBI (NCBI Release 102, released in 2018; (Thibaud-Nissen et al., 2013)). While long-read DNA sequencing technology was utilized to great effect to improve the reference *Harpegnathos* genome assembly, existing gene annotations still suffered from the shortcomings listed above, imposed by their reliance on traditional, short-read RNA-seq coupled with gene prediction software.

*Harpegnathos* is a promising model system to study the epigenetic regulation of brain and behavioral plasticity. Similar to colonies of other social insects, *Harpegnathos* colonies are founded by a mated reproductive female (“queen”) and contain many non-reproductive individuals (“workers”) that carry out all tasks necessary for colony survival. As in most social insect species (Keller and Genoud, 1997), *Harpegnathos* queens and workers differ greatly in reproductive physiology, social status and behavior, and lifespan, despite possessing the same genomic instructions. In addition, *Harpegnathos* ants display a rare form of phenotypic plasticity: workers retain the ability to convert to reproductive individuals called “gamergates” throughout their adult life (Bonasio, 2012; Peeters, 1991). In the absence of a dominant reproductive, *Harpegnathos* workers participate in a ritual dueling tournament, whereby winners become gamergates that activate their ovaries and acquire a queen-like social status (Peeters and Holldobler, 1995; Peeters et al., 2000). Workers that become gamergates lay eggs, cease activities associated with the worker caste (Gospocic et al., 2017), and acquire a longer lifespan (Ghaninia et al., 2017). Previous works identified changes in brain transcriptomes (Gospocic et al., 2017) and cell type proportions (Sheng et al., 2020) following this behavioral transition, establishing *Harpegnathos* as a powerful model for studying the epigenetic regulation of brain and behavioral plasticity.

Here, we use Iso-Seq long-read RNA sequencing to further improve the genomic infrastructure for genomic and epigenomic studies in *Harpegnathos* by generating more comprehensive annotations of splice isoforms and gene boundaries. These new annotations resulted in greatly improved analyses of bulk and single-cell RNA-seq, which revealed new caste-specific genes and splicing events and extended the reach of our single-cell atlas of the *Harpegnathos* brain.

## Results

### Using Iso-Seq to update *Harpegnathos* gene annotation

We previously generated a single-cell RNA-seq atlas of the *Harpegnathos* brain during the worker–gamergate transition and discovered extensive changes in cell type composition in glia and neurons (Sheng et al., 2020). While inspecting these sequencing data, we noticed that in many cases, even when using the latest NCBI annotation (NCBI Release 102; hereafter referred to as HSAL50), the single-cell RNA-seq reads mapped outside gene model boundaries, typically downstream of the annotated TTS of an example gene, *Ref1* (**Fig. 1A**, red box), resulting in decreased coverage and information loss. Motivated by examples such as this and by a desire to obtain a more comprehensive catalogue of splicing isoforms, we sought to improve the *Harpegnathos* gene annotations using PacBio Iso-Seq long RNA reads.

**Figure 1.**
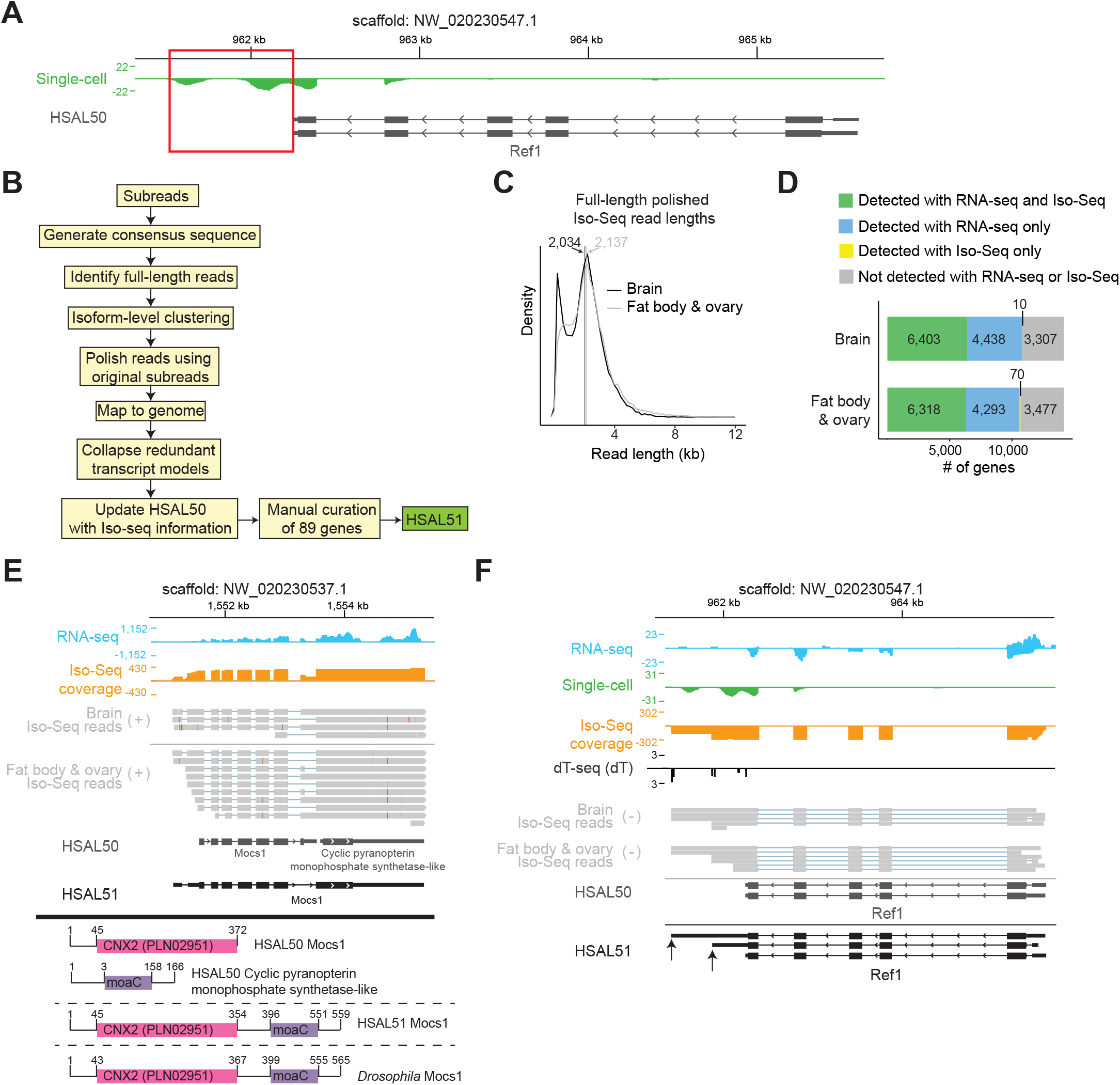
Iso-Seq improves gene models in *Harpegnathos*. (A) A gene with an incomplete 3’ UTR in the current annotation that precludes accurate quantification of single-cell RNA-seq signal. Pooled single-cell RNA-seq from worker (n = 6) and gamergate (n = 6) brains (Sheng et al., 2020) is shown. Scale represents counts per million. Red box indicates reads not assigned to the gene model. (B) Pipeline used for genome annotation. Iso-Seq reads from brains and fat body/ovary were processed and collapsed. The resulting Iso-Seq-based annotation was combined with the existing HSAL50 to produce an updated annotation, HSAL51. (C) Size distribution of polished, full-length Iso-Seq reads in brain and fat body/ovary. Vertical lines indicate median. (D) Number of genes detected in each Iso-Seq sample compared to short-read RNA-seq. (E) Example of a HSAL51 gene model derived from merging two incorrectly separated gene models in HSAL50. Coverage from short-read RNA-seq (blue) and Iso-Seq (orange) is shown in counts per million. Individual Iso-Seq reads from are also shown (gray). Below, the conserved domains on each relevant HSAL50 and HSAL51 gene are shown along with the conserved domains on *Drosophila Mocs1*. A subset of the HSAL51 *Mocs1* isoforms are shown. (F) Same locus as in (A), now including the updated HSAL51 annotation for *Ref1*, and showing RNA-seq and Iso-Seq coverage along with dT-seq and raw Iso-Seq reads. Scales represents counts per million. A subset of HSAL51 isoforms is shown and newly annotated TTS are indicated by arrows.

To maximize library complexity, we sequenced two separate polyA+ Iso-Seq libraries: one from a pool of *Harpegnathos* brains from different castes, and one from a mixture of ovary and fat body tissues. After processing the raw PacBio subreads (**Fig. S1A** and **Fig. 1B**), we obtained 34,867 and 33,520 full length “polished” reads with median length of 2,034 bp and 2,137 bp for the brain sample and the fat body/ovary sample, respectively (**Fig. 1C**). After aligning the polished Iso-Seq reads to the *Harpegnathos* genome, we compared gene coverage with that of previously obtained short-read RNA-seq in matching tissues (Shields et al., 2018). More than half of the genes detectable (RPKMs > 0.5) in our collection of deep short-read RNA-seq were also covered by Iso-Seq reads (**Fig. 1D**). As expected, genes detected by Iso-Seq tended to be more highly expressed (**Fig. S1B**). From the mapped reads, we collapsed redundant transcript models, generated predicted isoforms for each of the Iso-Seq libraries, and used these isoforms to refine the existing HSAL50 annotation. We further improved these models by manually adding 89 genes and reviewing all merged genes (**Fig. S1C** and **Table S1–3**) and designated this upgraded annotation as HSAL51.

Overall, HSAL51 contained 13,957 gene models that corresponded unambiguously to HSAL50 genes, 392 new genes either predicted from Iso-Seq signal or added manually (**Table S1–2**), and 57 gene models that merged two or more HSAL50 gene models into a single one in HSAL51 (**Fig. S1D** and **Table S3**). An example of a merged gene was the combination of two adjacent HSAL50 genes, each of which contained one of the two known *Drosophila melanogaster* MOCS1 protein domains; the resulting merged gene model for *Mocs1* in HSAL51 encodes a protein with a domain structure identical to its ortholog in *Drosophila* (**Fig. 1E**). Returning to the example of *Ref1* (**Fig. 1A**), Iso-Seq reads indicated the existence of at least two isoforms with TTS downstream of the one annotated in HSAL50, which captured the single-cell sequencing signal missed with the old annotation (**Fig. 1F**, arrows). For additional verification of the TTS predicted in our new annotation, we devised a custom RNA-seq protocol that compares short reads obtained with an anchored oligo-dT primer with random hexamers to remove signal from internal A-stretches and identifies the position of the polyA tail on mature mRNAs (“dT-seq”, **Fig. S1E–F**, see methods). In the case of *Ref1*, dT-seq signal analyses confirmed the existence of the new termination sites (**Fig. 1F**).

Thus, using long-read sequencing we updated the *Harpegnathos* gene annotation and recovered gene models that were split incorrectly in the HSAL50 assembly or that did not have a correctly annotated 3’ UTRs.

### Comprehensive annotation of transcriptional isoforms with Iso-Seq

Since its development in 2013 (Sharon et al., 2013), Iso-Seq has been performed on a genome-wide scale in a range of plants and animal species to improve the annotation of transcriptional isoforms (Abdel-Ghany et al., 2016; Beiki et al., 2019; Feng et al., 2019; Minio et al., 2019). The ability of Iso-Seq to sequence RNA molecules in their entirety confers an advantage in detecting splicing patterns compared to the typical short-read annotation strategy of relying on reads that cover a limited span across splice junctions. Indeed, HSAL51 contained a greater number of annotated transcripts with distinct splicing patterns (i.e. beyond simple extension of 5’ or 3’ UTRs) (**Fig. 2A**). In addition, gene models in HSAL51 exhibited more instances of all seven types of alternative splicing (Alamancos et al., 2015): skipped exon (SE), mutually exclusive exons (ME), alternative 5’ splice site (A5), alternative 3’ splice site (A3), retained introns (RI), alternative first exon (AF), and alternative last exon (AL) (**Fig. 2B**). Examples of genes with newly annotated alternative splicing events are presented in **Fig. S2A–C**.

**Figure 2.**
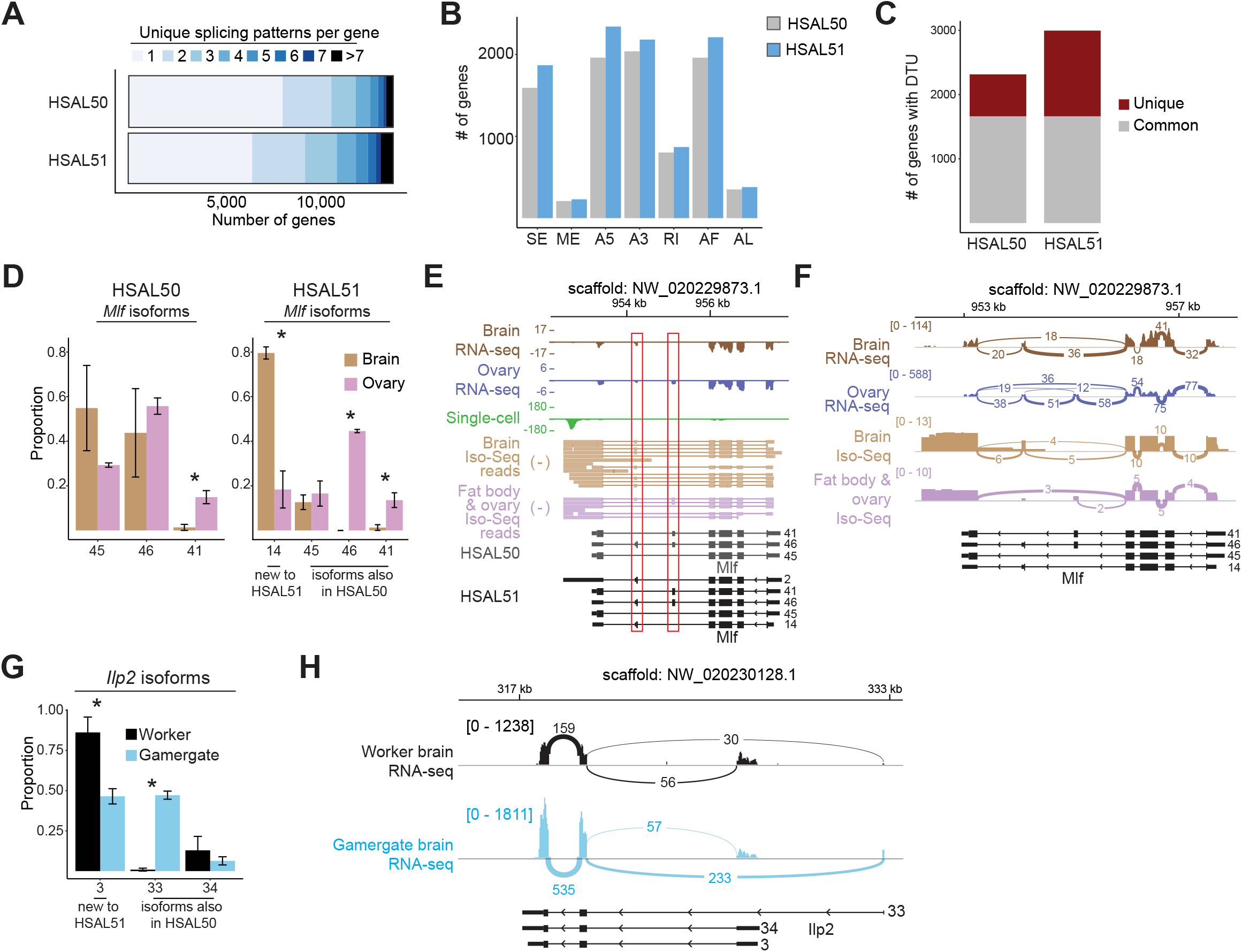
Differential transcript usage in tissues and castes. (A) Number of splicing isoforms per gene in HSAL50 and HSAL51. (B) Number of genes with select alternative splicing events (SE = skipped exon; ME = mutually exclusive exons; A5 = alternative 5’ splice site; A3 = alternative 3’ splice site; RI = retained intron; AF = alternative first exon; AL = alternative last exon) in HSAL50 and HSAL51. (C) Number of genes with differential transcript usage (padj < 10-5, maxDprop > 0.5, and rep_dtu_freq_threshold > 0.8) between at least two tissues out of brain, ovary, fat body, retina, optic lobe, and antenna. Genes identified as differentially spliced in both HSAL50 and HSAL51 are represented in gray, while genes identified in only one annotation are in red. (D) Tissue-specific isoform usage for *Mlf* (identified as DTU gene, padj < 10-10) according to HSAL50 (left) or HSAL51 (right) annotation. Proportions were calculated for each replicate, then averaged within condition. *, p < 0.05. (E–F) Genome browser (E) and sashimi plot (F) view of *Mlf* showing tissue-specific alternative isoforms. Splice junction line widths are scaled to the number of reads spanning the splice junction and the total number of reads mapped to *Mlf* for each tissue. (G) Caste-specific isoform usage in the brain for *Ilp2* (identified as DTU gene, padj < 10-10). Proportions were calculated for each replicate, then averaged within condition. *, p < 0.05. (H) Sashimi plot for the *Ilp2* gene. Splice junction line widths are scaled to the number of reads spanning the splice junction and the total number of reads mapped to *Ilp2* for each caste.

Next, we identified genes whose relative transcript expression varies between tissues (also called differential transcript usage, or DTU) (Van den Berge et al., 2019) using previously published bulk RNA-seq data from 6 *Harpegnathos* tissues: non-visual brain, ovary, fat body, retina, optic lobe, and antenna (Shields et al., 2018). Most genes (80%) exhibiting DTU in HSAL50 between at least two tissues were also detected in HSAL51, but the number of genes displaying DTU increased by 681 in the new annotation (**Fig. 2C**). For example, a newly annotated isoform (isoform 14) of the *myeloid leukemia factor* (*Mlf*) gene accounted for 80% of transcripts produced in the brain (**Fig. 2D**). This isoform contained exon 6 but not exon 5 of *Mlf* (**Fig. 2E-F**) and was not identified with short reads alone, highlighting the power of Iso-Seq in untangling complicated exon structures, especially in cases of mutually exclusive exons. Once annotated, short reads spanning the alternative splice junctions could be properly assigned to this isoform resulting in the identification of brain-specific DTU.

In addition to genes with DTU between tissues, we identified several genes with caste-specific transcript usage in the brain. One notable gene with caste-biased isoforms between workers and gamergates was *insulin-like-peptide 2* (*Ilp2*; *LOC105188195*), a gene similar to canonical insulin whose absolute transcript levels are higher in the brains or heads of reproductive individuals compared to those of non-reproductives in many ant species (Chandra et al., 2018). In general, insulin signaling has been identified as a key pathway regulating caste identity in social insects (Toth and Robinson, 2007). In addition to the higher gene-level *Ilp2* expression in brains of *Harpegnathos* gamergates compared to workers (**Fig. S2D**), isoforms 3 and 33 were used at different levels between the castes (**Fig. 2G**), with the upstream first exon being used more frequently in gamergates compared to workers (**Fig. 2H**). While these alternative splicing events appear to only affect the 5’ UTR, they might still have important consequences on insulin signaling, for example by regulating translation of the resulting mRNA, as observed in mammals (Minn et al., 2005; Shalev et al., 2002). Upon reanalysis of published data (Chandra et al., 2018; Patalano et al., 2015), two other ant species, the carpenter ant *Camponotus planatus* (Formicinae) and the giant ant *Dinoponera quadriceps* (Ponerinae) also displayed caste-biased selection of the first exon in brains (**Fig. S2E–F**). In both species, as in *Harpegnathos*, the reproductive caste was more likely than the non-reproductive caste to use the upstream first exon, suggesting that alternative splicing of *Ilp2* mRNA might be an evolutionary conserved mechanism for the caste-specific regulation of the insulin pathway.

Together, these analyses demonstrate that an Iso-Seq-enriched genomic annotation captures a greater complexity in the transcriptome which, in turn, can provide a comprehensive view of alternative splicing events between different biological samples—in this case tissues and castes.

### Extended 5’ and 3’ gene boundaries increase sensitivity of bulk RNA-seq

In addition to a more comprehensive view of transcriptional isoforms originating from alternative splicing, the long reads of Iso-Seq are expected to contain more complete UTRs in 3’ and, to some extent, 5’, resulting in more accurate annotation of TTS and TSS, respectively, extending the mappable regions of the gene models. Consistent with these expectations, for 45% of all gene models, the exons annotated in HSAL51 covered a larger (median extension, +20%) sequence than in HSAL50, whereas only 5% resulted in smaller gene models and even those were minimally impacted (median contraction, –2%) (**Fig. 3A–B**). This remarkable growth in annotated exonic space was largely due to the extension of 3’ UTR using newly annotated TTS (5,338 transcripts among 4,269 genes) and, to a smaller degree, to the extension of 5’ UTRs using newly annotated TSS (2,878 transcripts) (**Fig. 3C**). Transcripts were typically extended by more base pairs at the TTS compared to the TSS (**Fig. 3D**), with median extension length of 251 nt and 31 nt, respectively.

**Figure 3.**
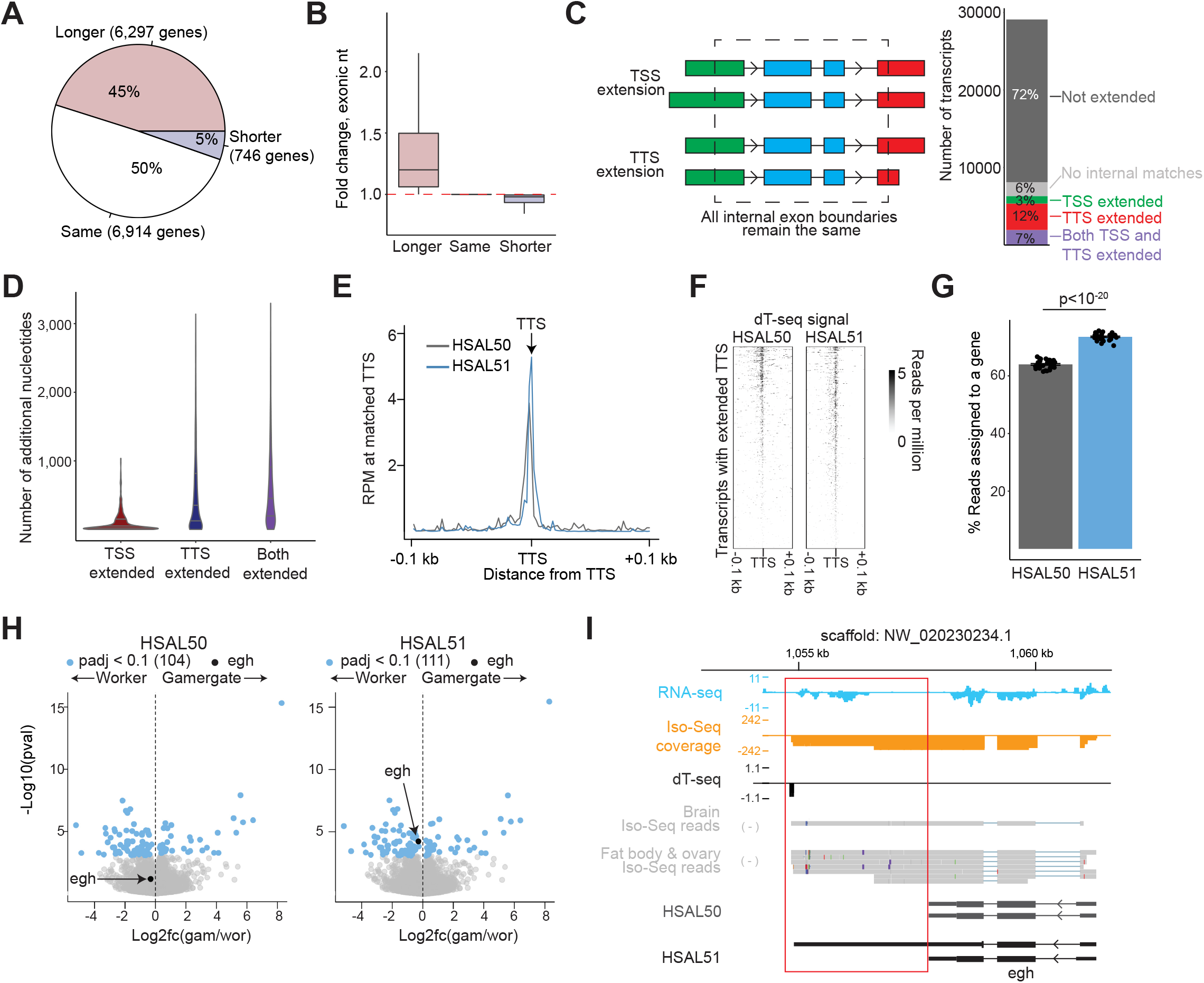
Extensions at 5’ and 3’ ends of genes improve analysis. (A) Number of genes whose exons cover the same, more, or fewer base pairs in HSAL51. (B) Fold-change in nucleotides covered by exons for each category in (A). (C) Scheme depicting TSS or TTS extension (left) and the percent of genes with no change, no match (no transcript with the same internal exon boundaries), TSS extension, TTS extension, or both TSS and TTS extension (right). (D) Average number of nucleotides added to transcripts with TSS and/or TTS extension. (E–F) Metaplot (E) and heatmap (F) of dT-seq coverage in a 0.2 kb window around TTS of transcripts with TTS extended by at least 200 nucleotides. (G) Percent of RNA-seq reads from worker brains mapping to features in HSAL50 or HSAL51. P-value is from a paired Student’s t-test. (H) Volcano plots of differential expression between gamergate (n = 12) and worker (n = 11) for HSAL50 (left) and HSAL51 (right). Genes with a p-adjusted < 0.1 are highlighted in blue. An example of a gene that is identified as differentially expressed in HSAL51 but not HSAL50 (*egh*) is highlighted in black. (I) Genome browser view of *egh* shows RNA-seq reads aligned to the extended portion of *egh*, which is supported by Iso-Seq reads. A subset of HSAL51 isoforms is shown.

To confirm that these 3’ UTR extensions originated from the annotation of *bona fide* TTS, we analyzed the position of polyA tails, as determined by dT-seq (see **Fig. S1E**), relative to the HSAL50 and HSAL51 gene models. As expected, the dT-seq signal accumulated at TTS in both annotations, and it was stronger at the TTS of HSAL51 gene models as compared to HSAL50 (**Fig. S3A**), including in a comparison of gene models with extended TTS in HSAL51 (**Fig. 3E–F**), demonstrating the accuracy of the newly annotated downstream TTS.

To evaluate the effect of the new annotation on RNA-seq mapping rates, we calculated the mapping rate of aligned reads using a newly generated dataset of *Harpegnathos* worker brains, which were not used in the construction of the HSAL50 (or HSAL51) annotations. Significantly more reads mapped to annotated exons in HSAL51 (**Fig. 3G**). This improvement in RNA-seq mapping rates had tangible benefits on the biological interpretation of sequencing datasets, such as, for example, the identification of additional differentially expressed genes in pairwise comparisons. Reanalyzing the worker *vs*. gamergate transcriptomes (Gospocic et al., 2017) with the new annotation identified the *egh* gene as significantly upregulated in workers (**Fig. 3H**). The *Drosophila* homolog of this gene is involved in the sex-peptide response and is implicated in the regulation of mating and egg-laying (Soller et al., 2006), suggesting a biological explanation for its caste-specific expression in *Harpegnathos*. Iso-Seq reads indicated the existence of a longer gene model for *egh* (**Fig. 3I**), which increased the number of reads mapping to this gene (**Fig. S3B**) and resulted in its confident identification as caste-specific.

These results show that the addition of Iso-Seq information extends the annotated gene models, allowing for the extraction of more information from RNA-seq experiments and resulting in higher sensitivity for differentially expressed genes.

### Iso-Seq-based annotation improves single-cell analyses

Given the 3’ bias of the most widely used techniques for drop-let-based single-cell sequencing, we reasoned that these analyses would be improved by the more accurate 3’ UTR annotations found in HSAL51. Indeed, the mapping rate of 10x Genomics single-cell RNA-seq reads from our previous worker–gamergates comparison (Sheng et al., 2020) increased in average by 44% (**Fig. 4A**). This resulted in substantially increased counts for a large majority of annotated genes, including several with important functions in the brain (**Fig. 4B**, below diagonal). In all 11 samples analyzed, increased mapping rates resulted in improvements for the total number of cells identified, as well as the average unique molecular identifiers (UMIs) and genes detected per cell (**Fig. 4C**). Overall, the total number of cells passing quality thresholds increased by 18% from 20,729 using the HSAL50 annotation to 24,560 using HSAL51 (**Fig. 4D, Fig. S4A**), showing the importance of accurate 3’ UTR annotations to extract the maximum amount of information from single-cell RNA-seq data.

**Figure 4.**
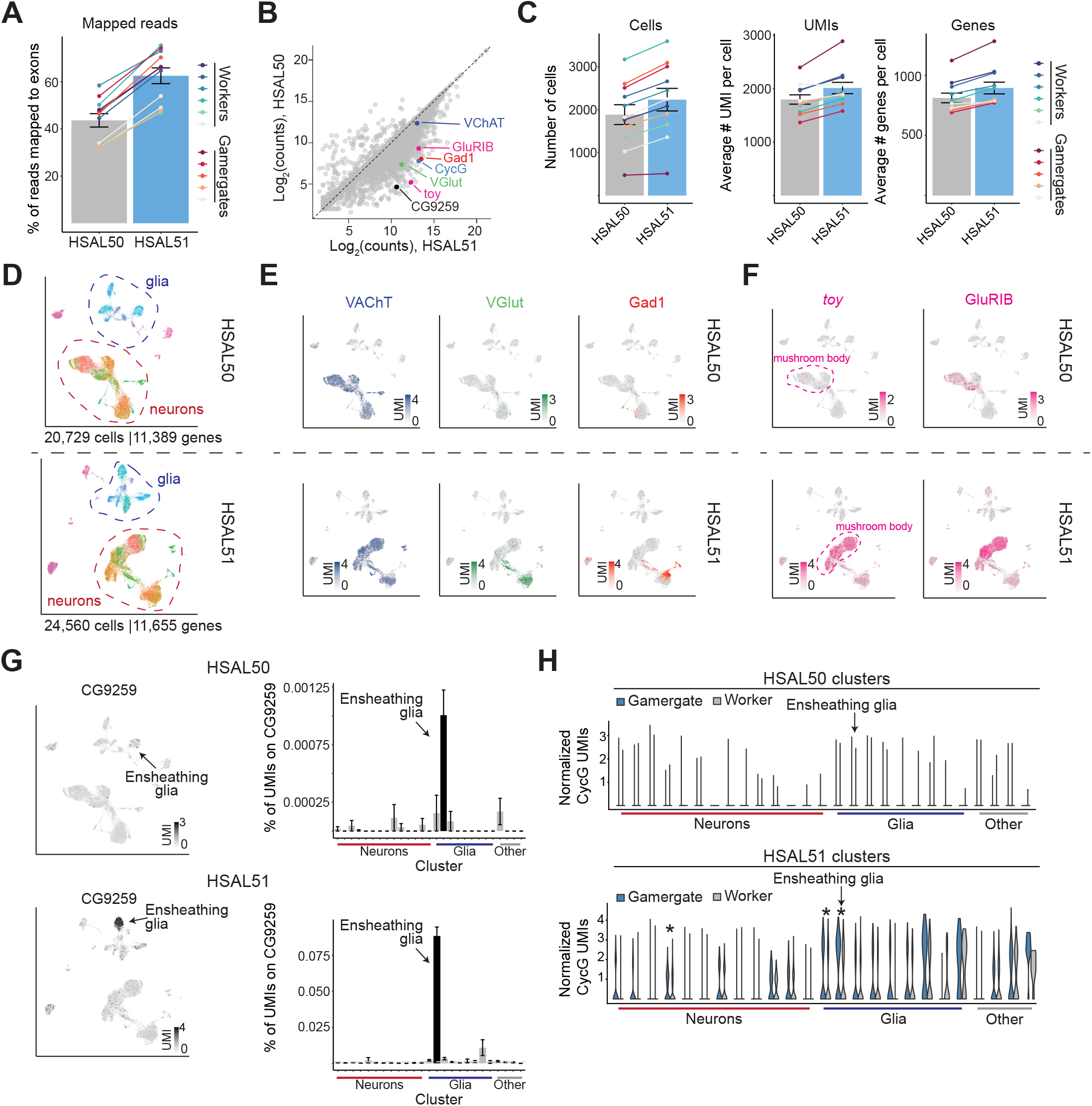
Iso-Seq assemblies improve single-cell sequencing analyses. (A) % of single-cell RNA-seq reads mapped to exons in HSAL50 vs. HSAL51. P-value is from a paired Student’s t-test. (B) UMI counts per gene in HSAL50 vs. HSAL51. Each dot is a gene. Genes of interest for subsequent analyses are highlighted. (C) Number of genes (left), mean number of UMI (center), and mean number of genes (right) using HSAL50 and HSAL51 annotations. (D) UMAP clustering of single-cell transcriptomes analyzed with HSAL50 or HSAL51 annotations. (E–F) Heatmaps showing normalized UMI counts for *VAChT, VGlut1*, and *Gad1* (E) or *toy* and *GluRIB* (F) in HSAL50 and HSAL51 analyses. (G) Heatmaps showing *CG9259* (left) and graph showing the % of UMIs mapping to *CG9259* in each cluster (right), in HSAL50 and HSAL51 annotations. (H) Violin plots for normalized UMIs for *CycG* in each cluster in HSAL50 and HSAL51 analyses, comparing cells in gamergate vs. worker brains. *, adjusted P-value < 0.05 by Wilcoxon-Rank Sum test with Bonferroni correction. All single-cell data were reanalyzed from (Sheng et al., 2020).

Markers for three major neuronal classes based on neurotransmitter usage, *VAChT* (cholinergic), *VGlut* (glutaminergic), and *Gad1* (GABAergic), were among the genes that benefitted from the increased mapping rates (**Fig. 4B**), resulting in clearer identification of cell types expressing these genes on the HSAL51 UMAPs (**Fig. 4E**). These improvements were not confined to genes associated with neurotransmitter usage; using HSAL51, we recovered in total 288 previously undetected marker genes with restricted, cell type-specific expression (**Fig. S4B**), likely missed in HSAL50 due to reads that were not assigned to the incomplete old gene models (**Fig. S4C**).

In addition, we recovered 12 new markers for mushroom body neurons (**Fig. S4D**), which are key to learning and memory in insects (Fahrbach, 2006; Strausfeld et al., 1998; Zars, 2000). Some of these markers were biased for mushroom body cells in HSAL50 but did not pass statistical thresholds due to overall low expression. Of these 12 newly identified mushroom body markers in *Harpegnathos*, 10 were previously described as mushroom body-specific genes in *Drosophila* (Davie et al., 2018; Jones et al., 2018) or honeybees (Traniello et al., 2020) (**Fig. S4D**). In particular, two *Harpegnathos* genes with homology to known mushroom body markers *GluR1B* and twin of eyeless (*toy*) (Crocker et al., 2016; Davie et al., 2018; Kurusu et al., 2000; Noveen et al., 2000), were barely detectable in HSAL50 but clearly mapped to mushroom body clusters in HSAL51 (**Fig. 4F** and **Fig. S4D**).

We previously showed that neuroprotective ensheathing glia cells are expanded during the worker–gamergate transition and lost at a faster rate in workers than in gamergates during aging (Sheng et al., 2020). Despite the in-depth investigation of this cell type in our previous study, the updated HSAL51 annotation allowed us to discover a new marker gene, *CG9259* (**Fig. 4G**), that was previously missed because the near-entirety of the single-cell RNA-seq reads fell within an extended 3’ UTR not annotated in HSAL50 (**Fig. S4E**). In *Drosophila, CG9259* encodes an ecdysteroid kinase-like protein (Aradska et al., 2015), suggesting that *Harpegnathos* ensheathing glia that express this gene might play an important role in the caste-specific regulation of the key developmental hormone ecdysone.

Finally, the increased single-cell transcriptome depth afforded by HSAL51 allowed us to identify differential gene expression between workers and gamergates within specific cell types, with more genes overall classified as caste-specific between single-cell clusters in the HSAL51 analysis (**Fig. S4F**). As with the newly detected marker genes, many of the 250 newly called differential genes had increased counts in HSAL51 compared to HSAL50, suggesting their new detection resulted from the increased mapping rates of the 3’-biased single-cell RNA-seq reads (**Fig. S4G**). For example, we identified *CycG* as a gene preferentially expressed in specific subtypes of gamergate cells as compared to their counterparts in workers (**Fig. 4H**). This observation is in agreement with previous studies reporting upregulation of *CycG* in the reproductive caste of various ant species (Nagel et al., 2020), including *Harpegnathos* gamergates (**Fig. S4H**). CycG regulates the insulin pathway (Fischer et al., 2015; Fischer et al., 2016), which, as mentioned above, is an important player in caste determination in social insects (Chandra et al., 2018; Toth and Robinson, 2007). The identification of the cell types that are the strongest drivers of CycG caste-biased expression will help inform future studies of this gene.

Thus, single-cell RNA-seq analyses were greatly improved by the increased accuracy of 3’ UTR annotations in HSAL51, resulting in 18% more single cells identified computationally, a clustering of transcriptional types more reflective of biological function, recovery additional cell-type markers, and higher sensitivity for differentially expressed genes.

### Long non-coding RNAs in single-cell sequencing analysis revealed by Iso-Seq

Protein-coding genes are often the focus of transcriptomic studies, but many genes are transcribed into noncoding RNAs with important regulatory roles (Bonasio and Shiekhattar, 2014; Rinn and Chang, 2012; Shields et al., 2019). Similar to the case of protein-coding transcripts, several gene models for various types of noncoding RNAs, and in particular long noncoding RNAs (lncRNAs) were also extended in HSAL51 compared to HSAL50 (**Fig. 5A, Fig. S5A**), although not to the same extent, possibly due to their overall lower expression level.

**Figure 5.**
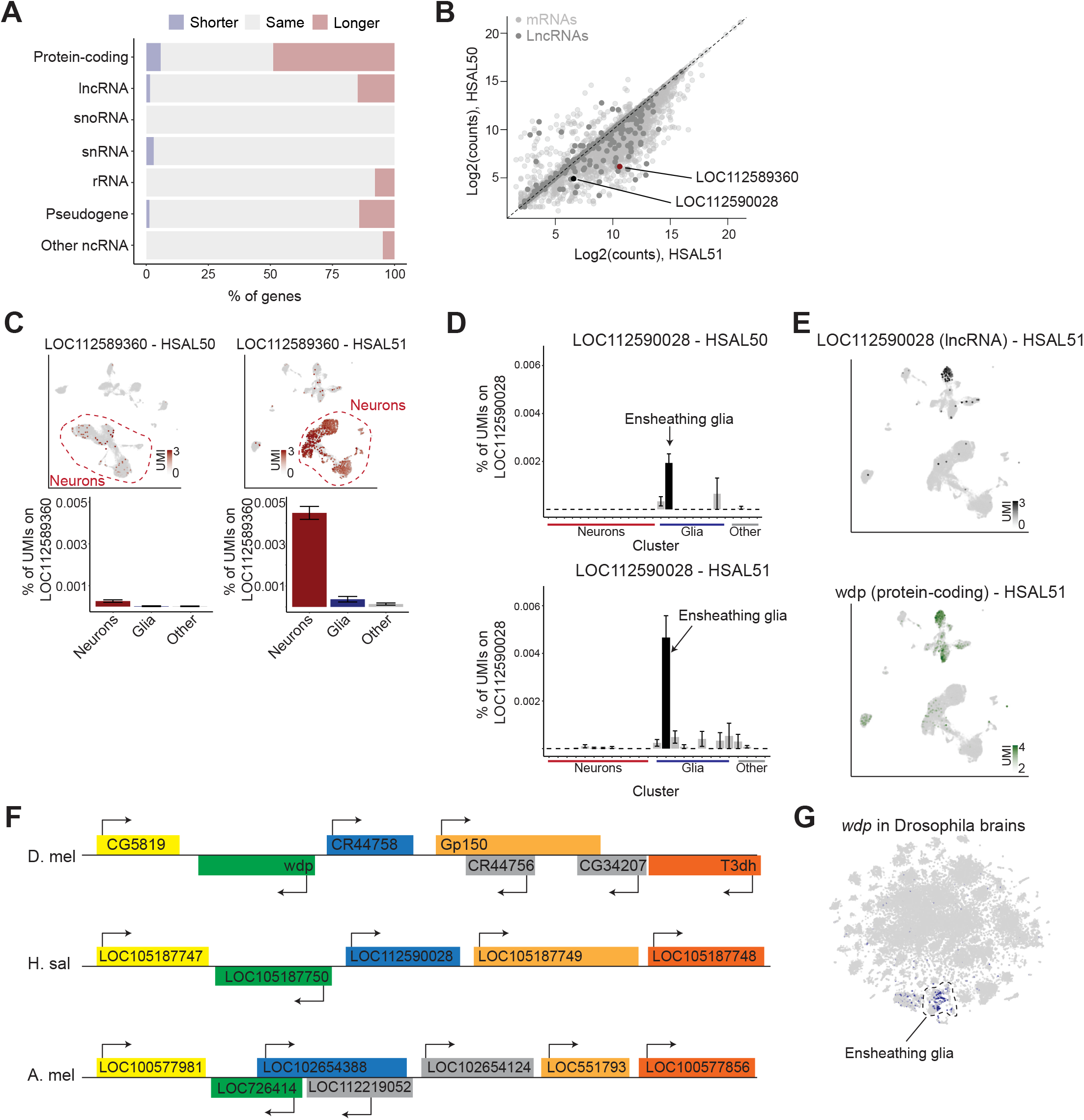
Single-cell characterization of lncRNA expression in the *Harpegnathos* brain. (A) Number of genes whose exons cover the same, more, or fewer base pairs in HSAL51 compared to HSAL50, classified by biotype. (B) Scatter plot for UMI counts per gene in HSAL50 vs. HSAL51. LncRNAs are highlighted in dark gray, and gene of interest mentioned in subsequent panels are shown in black or red. (C) Heatmap showing normalized UMI counts of neuron-specific lncRNA *LOC112589360* in HSAL50 and HSAL51 (top) and % of UMIs mapping to *LOC11258930* in each cell type (bottom). (D) % of UMIs mapping to ensheathing glia-specific lncRNA *LOC112590028* in HSAL50 (top) and HSAL51 (bottom) (E) Heatmap with normalized UMI counts of lncRNA *LOC1125890028* (top) and protein-coding gene wdp (bottom). (F) Diagram showing relative positions of genes in the vicinity of *wdp* in *Drosophila, Harpegnathos*, and *Apis mellifera*. Orthologous genes are the same color. (G) Heatmap showing normalized UMI counts of *wdp* in single-cell RNA-seq from *Drosophila* brains (Davie et al., 2018). All single-cell data were reanalyzed from (Sheng et al., 2020).

Single-cell analyses using the updated gene models in HSAL51 revealed 130 lncRNAs with more UMIs compared to HSAL50 (**Fig. 5B**), suggesting that the new annotation might provide additional insight on the patterns of lncRNA expression in the *Harpegnathos* brain. We recovered a set of lncRNAs with neuronal-specific expression profiles, some of which could not be detected using HSAL50 gene models (**Fig. S5B**). One example was LOC112589360, which had over 20 times mapping reads in HSAL51 compared to HSAL50 (**Fig. 5B**), with a corresponding increase in its calculated expression levels in neurons (**Fig. 5C**), as well as the fraction of neurons where this lncRNA could be detected, from 0.6% in HSAL50 to 10.1% in HSAL51 (**Fig. S5B**).

In addition to these neuronal lncRNAs, we identified several cell type-specific lncRNAs in ensheathing glia. LOC112588339. LOC112588340 was merged from two adjacent lncRNAs annotated in HSAL50, with Iso-Seq reads clearly supporting the HSAL51 gene model (**Fig. S5C**). The new merged gene model had negative coding potential as assessed by CPC and PhyloCSF (Kong et al., 2007; Lin et al., 2011) and was one of the strongest markers of ensheathing glia (**Fig. S5D**). Another updated lncRNA, LOC112590028, was specific to ensheathing glia, but missed by previous analyses due to low mapping rates in HSAL50 (**Fig. 5D-E**). The protein-coding gene adjacent to this lncRNA, *windpipe* (*wdp*), was also preferentially expressed in ensheathing glia (**Fig. 5E** and **Fig. S5E**), suggesting potential co-regulation of the coding and non-coding transcript in *cis* as previously reported for other lncRNAs-mRNA pairs (Engreitz et al., 2016). *Drosophila wdp* is a transmembrane protein with known functions in the wing disc (Takemura et al., 2020) and the trachea (Huff et al., 2002), and it has also been implicated in synaptic target recognition (Kurusu et al., 2008) and learning (Williams-Simon et al., 2019). While the sequence of the lncRNA LOC112590028 itself is not conserved, *Drosophila* has a lncRNA, *CR44758* in the same position as LOC112590028, between *wdp* and *Gp150* (**Fig. 5F**), suggesting that synteny of this locus, and potentially its molecular regulation, have been maintained over 350 million years of divergent evolution (**Fig. 5F**). In fact, *wdp* is also expressed specifically in *Drosophila* ensheathing glia, further supporting a conserved regulation of this locus across distantly related insect species (**Fig. 5G**).

Thus, similar to protein-coding genes, lncRNA annotations were also improved by the addition of Iso-Seq data and this resulted in increased visibility of these regulatory transcripts in single-cell analyses.

## Discussion

Genomic resources are becoming increasingly common for a wide range of species, beyond traditional model organisms, facilitating molecular analyses of an ever-growing variety of biological phenomena. However, in addition to high-quality genome assemblies, accurate gene annotations are indispensable for genome-wide studies. Here, we used Iso-Seq to more comprehensively annotate genes in *Harpegnathos* and showed that these improved annotations result more accurate detection of alternative splicing events, increased sensitivity of differential gene expression analyses, and deeper single-cell transcriptomes.

### Long RNA reads for more complete gene annotations

Often, genome annotations are constructed *de novo* from short-read RNA-seq, with or without the guide of an existing genome assembly (Haas et al., 2013; Haas and Zody, 2010; Martin and Wang, 2011). While RNA-seq-based annotations are typically sufficient to identify protein-coding mRNAs and characterize their expression levels in bulk RNA samples, they are limited in their ability to fully annotate complete transcript isoforms, extended UTRs, and lncRNAs. Because of its ability to sequence long RNA molecules in a single read, Iso-Seq overcomes these limitations and has already been used to identify novel transcriptional isoforms in various genomes (Abdel-Ghany et al., 2016; Wang et al., 2016) including the very well annotated human genome (Sharon et al., 2013). Ideally, these more comprehensive isoform maps can be used to detect genes with differential splicing between biological conditions. In human cells, targeted long-read sequencing was used to examine splicing of neurexins (Treutlein et al., 2014), leading to the association of aberrant splicing of NRXN1 with psychosis disorders (Flaherty et al., 2019).

Incomplete annotations can impede proper analysis of massively high-throughput single-cell RNA-seq, most of which is heavily biased towards the 3’ end of the gene due to oligo-dT capture by beads (Zhang et al., 2019). Thus, having a correct annotation of the 3’ UTR and the TTS of genes becomes crucial. Annotation of the 3’ ends of genes is hampered by widespread occurrences of multiple polyadenylation and cleavage sites (Beaudoing and Gautheret, 2001; Wang et al., 2008) and intrinsic limitations of existing annotation methods (Shenker et al., 2015). Even in very high-quality reference genomes, such as the human genome, reads from single-cell RNA-seq can fall past the annotated 3’ UTR, resulting in information loss (Ntranos et al., 2019). Previous studies have employed computational strategies to improve the assignment of these reads to gene models, including extending every 3’ UTR by 2 kb (Sebe-Pedros et al., 2018) or mapping to non-overlapping windows on the genome and assigning each window to a gene based on the proximity to an annotated TTS (Sa et al., 2020). Although methods exist that model 3’ UTR annotations based on deviation in RNA-seq coverage (Huang and Teeling, 2017; Shenker et al., 2015), here we employed that an empirical approach, based on long read sequencing, to improve the annotation of 3’ UTRs (**Fig. 1**).

### Splicing analysis with additional Iso-Seq information

Differential alternative splicing occurs between tissues and biological conditions (Baralle and Giudice, 2017; Streuli and Saito, 1989; Xu et al., 2002). Specific isoform usage, potentially mediated by varying RNA-binding protein expression, is widespread and affects protein-protein interaction networks (Buljan et al., 2012; Ellis et al., 2012; Wang et al., 2008). Identification of novel isoforms with Iso-Seq revealed genes with differential splicing patterns between tissues, including one gene with a newly annotated mutually exclusive exon (**Fig. 2**).

Caste-specific alternative splicing has been observed in social insects. Doublesex (*dsx*) is differentially spliced between queens and workers in the ants *V. emeryi, S. invicta*, and *W. auropunctata* in addition to its differential splicing in non-social and social insects between males and females (Burtis and Baker, 1989; Cho et al., 2007; Mine et al., 2017; Miyakawa et al., 2018). While we did not find splicing changes between worker and gamergate in *dsx*, we identified caste-specific isoform usage in the brain for *Ilp2*, a known factor in caste determination that is also differentially expressed on the gene-level in the brains of many ant species including *Harpegnathos* (Chandra et al., 2018). Other ants also seem to have caste-biased splicing of the first exon, similar to *Harpegnathos*, suggesting a possible conserved mechanism that will require more investigation to understand.

### Bulk and single-cell RNA-seq analysis with refined gene models

The integration of Iso-Seq into the annotation resulted in improved gene models mostly due to an extension of first and last exon, corresponding to the 5’ and 3’ UTR, respectively (**Fig. 3A– F**). Many 3’ UTR extensions were confirmed by the presence of dT-seq signal, designed to capture the location of non-templated polyA tails. The new annotation captured more information from RNA-seq, with a median of 15% more reads. Increased mapping rates had immediate tangible effects on discovery, as a reanalysis of existing RNA-seq data from the worker–gamergate transition led to the identification of 7 new caste-specific genes in the brain. One of these, *egh*, was previously implicated in reproductive behavior in *Drosophila* (Soller et al., 2006), and therefore represents a high value candidate for the dissection of the molecular regulation of social behavior in ants.

The improvements for single-cell RNA-seq (**Fig. 4**) were even more remarkable. Mapping existing single-cell RNA-seq reads from gamergate and worker brains to the new annotation improved the median percent of reads mapped to exons from 47% to 66%. The increased mapping resulted in 18% more cells passing the minimum UMI and gene thresholds and the addition of 266 new cell type-specific genes to our single-cell atlas of the *Harpegnathos* brain. The extraction of more information from this existing dataset provided several new biological insights. A number of biologically relevant genes had increased UMI mapping in the new HSAL51 annotation that translated to improved identification and visualization of cells expressing these genes, including established mushroom body markers *toy* and *GluIRB*, and the neurotransmitter-associated markers *VAChT, VGlut*, and *Gad1*. Using the new annotation, more genes were classified as specific to the mushroom body or to a specific cell type. We identified *CG9259* as a new protein-coding gene specifically expressed in *Harpegnathos* ensheathing glia, a cell type previously linked to caste regulation and aging (Sheng et al., 2020).

The new annotation also improved analysis of lncRNAs in the single-cell data set (**Fig. 5**) revealing a lncRNA that is a marker of these ensheathing glia and is expressed in a similar set of cells as its adjacent protein-coding gene *wdp*. In single-cell analysis of the *Drosophila* brain, *wdp* expression is largely restricted to ensheathing glia, though detection of the lncRNA syntenic to the *Harpegnathos* ensheathing glia marker LOC112590028 is not detected in this data set. Further work would be required to confirm the link between LOC112590028 and *wdp* expression in *Harpegnathos*, but *wdp* expression patterns in *Drosophila* suggest a conservation in the regulation and, possibly, function of this gene.

### Conclusions and outlook

The new *Harpegnathos* annotation, bolstered by Iso-Seq and new RNA-seq, allowed us to uncover new patterns of differential alternative splicing, differentially expressed genes, and markers of cell types in single-cell sequencing. The identification of new candidate genes involved in tissue-specific and caste-specific regulation of gene expression highlights the advantages of a more complete gene annotation. Future genomic studies in *Harpegnathos* will undoubtedly benefit from these improved annotations, which will help obtain new insights on the molecular regulation of their remarkable phenotypic plasticity.

Beyond the implications for *Harpegnathos* genomics, our analyses show that Iso-Seq is an effective strategy for improving incomplete gene annotations, maximizing the amount of information garnered from genome-wide sequencing data.

## Materials and Methods

### Ant colonies and husbandry

As previously described (Sheng et al., 2020), *Harpegnathos* ants were descended from a gamergate colony collected in Karnataka, India in 1999 and bred in various laboratories. Ant colonies were housed in plaster nests in a clean, temperature-and humidity-controlled ant facility on a 12 hour light/dark cycle. Ants were fed three times per week with live crickets.

### PacBio Iso-seq

Non-visual brains and combined samples of fat body and ovary were dissected from a variety of castes and aged *Harpegnathos* ants. RNA was purified with TRIzol and sequenced on the PacBio Sequel II on one 8M chip at the University of Washington PacBio sequencing center.

### RNA-seq of *Harpegnathos* worker brains (Fig. 3G)

Brains from transitioning *Harpegnathos* ants were dissected out in phosphate-buffered saline, placed in TRIzol (Invitrogen #15596026) and stored at -70 C until RNA extraction. Each brain was processed separately. To extract RNA, thawed brains were homogenized in TRIzol by pestle, and frozen and thawed again. Chloroform was added, followed by vigorous vortexing and centrifugation at 21,000 g for 10–15 minutes at 4 C. The aqueous phase was purified using RNA Clean and Concentrator kit (Zymo Research #R1013) following manufacturer’s instructions. Extracted RNA was quantified using NanoDrop 2000 (Thermo Scientific) and RNA integrity was checked using High Sensitivity RNA ScreenTape (Agilent #5067-5579, 5067-5580, 5067-5581)

on a TapeStation 2200 or 4200 (Aglient). 250–500 ng extracted RNA was used to prepare libraries with NEBNext Ultra II Directional RNA Library Prep Kit for Illumina (New England Biolabs #E7760S) following manufacturer’s instructions. Libraries were quantified using Qubit dsDNA High Sensitivity Assay Kit (Invitrogen #Q32854) on a Qubit 2.0 fluorometer (Invitrogen) and fragment size distribution was checked using High Sensitivity D1000 ScreenTape (Agilent #5067-5584, 5067-5585, 5067-5587) on a TapeStation 2200 or 4200 (Aglient). Libraries were combined in two pools and sequenced in two runs of a NextSeq 500 machine (Illumina) in HighOutput mode with a 2x 150 bp configuration.

### Iso-seq data processing and annotation construction

Full-length Iso-seq reads were classified from circular consensus sequences using lima (Pacific Biosciences; https://github.com/PacificBiosciences/barcoding) and refined with isoseq3 (Pacific Biosciences; https://github.com/PacificBiosciences/IsoSeq) with --require-polya to filter for reads ending in a polyA tail. Fulllength reads (FLNC) were clustered using isoseq3 cluster and polished using isoseq3 polish. FASTQ files output from polishing step were mapped using the STARlong module of STAR (Dobin et al., 2013) to the *Harpegnathos* genome (GCF_003227715.1) with parameters suggested for mapping Iso-Seq reads to a genome provide by cDNA_Cupcake (https://github.com/Magdoll/cDNA_Cupcake/wiki/Best-practice-for-aligning-Iso-Seq-to-ref-erence-genome:-minimap2,-deSALT,-GMAP,-STAR,-BLAT). Redundant transcript models were collapsed using TAMA Collapse (Kuo et al., 2020). Transcript models were generated separately for brain and fat body/ovary tissues, and were merged together with the existing annotation produced by NCBI (GCF_003227715.1) using TAMA Merge (Kuo et al., 2020), with the no_cap option and prioritizing the brain Iso-Seq models followed by the fat body/ovary Iso-Seq models and then the existing annotation.

Several automatic and manual processing steps were performed to refine this annotation (see **Fig. S1C** for an overview). The mitochondrial scaffold and odorant receptor were annotation manually due to the challenges of annotating these genes computationally (see below, “Manual annotation of mitochondrial scaffold and odorant receptor genes”). A list of the odorant genes added and the genes from the previous annotation that were replaced can be found in **Table S1**. Genes “merged” in the Iso-Seq annotation (evidence suggests that two gene models should be one gene model; see example in **Fig. 1E**) were manually reviewed, with evidence from Iso-Seq, bulk RNA-seq, and homology of the genes in question taken into account to ensure that genes were not spuriously merged.

Due to suspicions of transcript models with retained introns representing pre-mRNAs sequenced by Iso-Seq, new transcripts with retained introns found by SUPPA (Alamancos et al., 2015) were subjected to further filtering. BLASTn was used to find homologs for transcript models with and without the retained intron, using a BLAST index created from all transcripts from *Drosophila melanogaster* (GCF_000001214.4), *Apis mellifera* (GCF_003254395.2), *Nasonia vitripennis* (GCF_009193385.2), and *Bombyx mori* (GCF_014905235.1) NCBI annotations. The query cover for the pairs of transcripts with or without retained introns were compared. If the transcript model without the retained intron had higher query coverage, the transcript with the retained intron was removed from the annotation.

From these final gene and transcript models, Transdecoder (Haas and Papanicolaou) was run to find coding sequences. The longest ORFs were BLASTed (BLASTp) to a reference of proteins from *Drosophila melanogaster, Apis mellifera*, and *Homo sapiens* (GCF_000001405.39) with an e-value cutoff of 1e-5.

### Manual annotation of mitochondrial scaffold and odorant receptor genes

The scaffold NW_020230424.1 was identified as the mitochondrial scaffold, as it contained the best BLAST hits of *Drosophila* mitochondrial genes. Several single-copy *Drosophila* genes had multiple hits on this scaffold and many genes were annotated as pseudogenes, necessitating a manual reannotation of these loci. We performed BLAST of the NW_020230424.1 sequence against itself and established that the positive strand of the 3’ end of the scaffold aligns with the positive strand of its 5’ end. This alignment pattern and the double BLAST hits of the *D. melanogaster* genes are consistent with NW_020230424.1 being an erroneous linear assembly of ∼2 iterations of the circular mtDNA sequence. Furthermore, visual examination of aligned RNA-seq reads revealed an abnormally high number of base mismatches and small indels in this scaffold. This suggests that NW_020230424.1 may have not been optimally polished when the genome was being assembled and explains why most genes contain frame shifts and/ or premature stop codons and are annotated as pseudogenes.

Considering these assembly and annotation issues, we removed existing gene models in NW_020230424.1 and manually re-annotated coding genes in its 3’ half. We identified regions that exhibited contiguous coverage in bulk RNA-seq, looked for predicted ORFs in such regions, and performed BLAST of their translations against *Drosophila* mitochondrial proteins. We also took into account gene order, as the mitochondrial genomes of *Drosophila* and *Harpegnathos* are completely syntenic (Garesse, 1988). In the instances where putative assembly errors caused frame shifts and/or premature stops, we split the genes into several fragments and added an alphabetical index to the fragments’ names. For example, *Drosophila mt:Cyt-b* corresponds to *mt:-Cyt-ba, mt:Cyt-bb*, and *mt:Cyt-bc* in *Harpegnathos*.

For the odorant receptors, we converted annotations manually generated for previous versions of the *Harpegnathos* genome (Zhou et al., 2012) the current genome assembly. We mapped the mRNAs of these old predictions to the current genome assembly using exonerate v. 2.2.0 and visually compared them to the HSAL50 annotations of the corresponding loci (Slater and Birney, 2005). Where necessary, we manually updated HSAL51 predictions taking into account the exon-intron structure of the mapped annotations of Zhou et al. and bulk RNA-seq coverage.

### dT-sequencing

Using previously isolated RNA (Shields et al., 2018) from worker tissues (fat body, ovary, non-visual brain, and optic lobe) and gamergate tissues (fat body, ovary, and non-visual brain), a version of RNA-seq was performed to specifically capture 3’ ends of transcripts with a polyA tail, here called “dT-seq” (see **Fig. S1E** for overview). RNA was fragmented by adding 4 µL of 5x 1st strand buffer from SSIII RT kit (Invitrogen catalog #18080-044) to RNA and incubating at 94°C for 16 minutes, followed by a ramp down to 4°C. PolyA+ selection was performed with OligodT25 dynabeads (Invitrogen catalog #610-02) with 3 washes with Oligo-dT washing buffer and eluted with BTE (10 mM bis-tris pH 6.7, 1 mM EDTA) containing either oligo-dT primer or random hexamer primers. Library construction was completed following established protocols (Shields et al., 2018) in parallel for the oligo-dT-primed and the random hexamer-primed samples. Libraries were sequenced in paired-end mode on a NextSeq500.

### dT-sequencing analysis

FASTQs from oligo-dT primed samples (“dT”) were filtered to keep only reads with at least 5 Ts with one mismatch at the 5’ end of the read, then trimmed using prinseq (Schmieder and Edwards, 2011) to remove tails from reads. These reads and random hexamer reads (“hex”) were aligned to the *Harpegnathos* genome using STAR with default parameters, except --alignIntronMax 50000.

Hex samples were used to filter out reads coming from polyA tracts within a transcript (see **Fig. S1E**, bottom). Genomic coverage of the first read (pair closest to polyA tail) was computed for both dT and hex samples using GenomicRanges (Lawrence et al., 2013). For each “peak,” defined as a contiguous region of coverage with at least one read, tentative summits were defined as any position with coverage at least 90% of the highest read total within the peak. The “summit” of each peak was defined as the most downstream tentative summit. dT peaks were sorted into three categories: (1) dT peaks that did not overlap with a hex peak, (2) dT peaks whose peak was upstream of a hex peak – see **Fig. S1E**, “discarded” box (3) dT peaks whose summit was downstream or equal to the hex summit – see non-discarded peaks in **Fig. S1E**. Peaks in categories (1) and (2) were discarded, leaving a list of dT peaks that did not have hex signal downstream, indicating dT reads coming from polyA tails instead of internal polyA stretches. Reads overlapping these peaks were retained and used for further analysis and for any genome browser snapshots showing dT signal.

To verify that the retained dT reads were from the end of transcripts (see **Fig. S1F**), aligned dT and hex reads were reduced to the first base at the 5’ end of the read using GenomicRanges. dT “peaks” were again detected using the strategy above and categorized as 3’ UTR peaks or CDS peaks based on their position within HSAL51 transcripts. Read coverage for dT and hex reads were computed at each of these peak sets.

### Splicing

Isoform-level counts from *Harpegnathos* samples were generated using kallisto (Bray et al., 2016) with HSAL50 or HSAL51 annotations with any single-exon transcripts removed. For gamergate and worker brains (Gospocic et al., 2017) (single-end sequencing), kallisto quant was run with the parameters -b 30 --rf-strand-ed--single -l 200 -s 1. For tissue samples (non-visual brain, ovary, fat body, antenna, retina, optic lobe; paired-end) (Shields et al., 2018), kallisto quant was run with the parameters -b 30 --rf-stranded. Differential transcriptional usage was tested with RATS (Froussios et al., 2019) with the parameters p_thresh=0.05, dprop_thresh=0.2, abund_thres=5 (for tissues) or p_thres=0.01,d-prop_thresh=0.1, abund_thresh=1 (for gamergate/worker brains). The number of genes with DTU using each annotation (**Fig. 2C**) was the number of gene with a padj<1e^-5^, maxDprop>0.5, and rep_dtu_freq_threshold>0.8 between any two tissues. Proportions of transcript usage was calculated for each replicate using kallisto TPMs. Proportions for each isoform were calculated by averaging replicates.

### Analysis of extended transcripts and genes

Genes with additional exonic nts were identified by comparing the total number of nts covered by exons of the gene in HSAL50 and HSAL51.

Genes with extensions of their transcription start or termination sites were identified by first finding pairs of HSAL50 transcripts with a corresponding HSAL51 transcript with the same internal exon structures in the two annotations (see **Fig. 3C**, left), defined by transcripts with all exon boundaries the same except for the start and/or termination sites. A small number of HSAL50 transcripts had no matching transcript in the HSAL51 annotation (light gray, “no internal matches” in **Fig. 3C**). The start and termination sites of the remaining HSAL50 transcripts were compared to their paired HSAL51 transcripts and sorted into categories of not extended, TSS extended, TTS extended, and both TSS/TTS extended.

### Bulk RNA-seq analysis

Bulk RNA-seq data for *Harpegnathos* from RNA newly sequenced from worker brains (see above) or previously published worker/ gamergate brains (Gospocic et al., 2017) or tissues (Shields et al., 2018), was analyzed using STAR with default parameters except --alignIntronMax=50000. Previously published RNA-seq from brains of *Camponotus planatus* (PRJNA472392) (Chandra et al., 2018) and *Dinoponera quadriceps* (PRJNA255520) (Patalano et al., 2015) was aligned with the same parameters to the *Camponotus floridanus* (GCF_003227725.1) and *Dinoponera quadriceps* (GCF_001313825.1) genomes and annotations, respectively, as no genome assembly or annotation has been published for *Camponotus planatus*. Read counts or TPM (**Fig. S1B, Fig. S2D** and **Fig. S4H**) were produced by an in-house script using GenomicRanges summarizeOverlaps (counting mode = union) (Lawrence et al., 2013) that counts the number of reads overlapping each gene model. Differential expression analysis was performed using DESeq2 (Love et al., 2014).

Single-cell tracks shown as examples were produced by aligning previously published single-cell RNA sequencing from *Harpegnathos* workers and gamergates (Sheng et al., 2020) to the *Harpegnathos* genome aligned using STAR with default parameters except --alignIntronMax=50000 (Dobin et al., 2013).

### Genome browser screenshots

All genome browser screenshots were produced using IGV v2.8.6. Bigwig tracks for visualization were produced using DeepTools (Ramirez et al., 2016). Sashimi plots were produced using IGV v2.8.6, with a custom scaling of splicing lines (line widths scaled to the total number of reads mapped to the locus, with a constant scaling factor used between sequencing from different castes for each technology).

### Single-cell analysis

Single-cell RNA sequencing from brains of *Harpegnathos* workers and gamergates previously generated with 10X Genomics (Sheng et al., 2020) was reanalyzed using CellRanger (Zheng et al., 2017) with default parameters and either the HSAL50 or HSAL51 annotations provided. CellRanger was used to produce digital gene expression matrices for all samples for both annotations. These matrices were processed by Seurat v3 (Butler et al., 2018). Cells with at least 200 genes and 500 UMIs were retained, with genes required to be expressed in at least 3 cells. UMIs were log-normalized with a scale factor of 10,000, the top 2,000 variable features detected using the “vst” selection method, and data were scaled so the mean of each gene across cells was 0 and variance was 1. As the samples were produced in three separate batches, the experiment was regressed out during this step. Cells were clustered using the variable features previously detected.

Clustering and visualization were performed with Seurat’s principal component analysis followed by JackStraw to detect significant principal components. All components were selected until a component had a P > 0.05. Cells were clustered with a resolution of 1 and clusters were visualized using UMAP. Cell type for each cluster was performed using previously established markers (Sheng et al., 2020).

For cluster-level pseudobulk expression analyses (“% UMIs on gene”), the number of UMI for each gene from all cells in each cluster for each sample were added together and normalized by the total number of UMI detected in that sample and cluster.

To compare clusters from HSAL50 and HSAL51, cluster groupings were produced by comparing the cells present in each cluster, which largely stayed constant between analyses using the two annotations. In all cases, one cluster from HSAL51 corresponded to one or two clusters from HSAL50 (in two cases, one HSAL51 cluster was split into two clusters in HSAL50).

Marker genes for each of these cluster groups (**Fig. S4B** and **C**) were defined using Seurat FindAllMarkers with the parameters only.pos=TRUE, min.pct=0.25, logfc.threshold=1. Only markers with a p-adjusted less than 0.05 were retained. Differentially expressed genes between castes within each cluster were identified using Seurat FindMarkers, run within each cluster with worker cells and gamergate cells provided as the two identities and default parameters. Only genes with a p-adjusted less than 0.01 were retained.

### Mushroom body markers

For **Fig. S4D**, mushroom markers in the HSAL50 and HSAL51 analysis were found using FindMarkers with the mushroom body clusters (defined by *mub*) as one group and all other cells as another group, with the parameters only.pos=T, min.pct=0.25, and logfc.threshold=1.5. Markers detected in HSAL51 and not HSAL50 were classified as new mushroom body markers and heatmaps of their expression in cells in each cluster was plotted using Seurat DoHeatmap. Lists of mushroom body enriched genes found by comparing *Drosophila* head to mushroom body transcriptomes (“D. mel MB vs head”) (Jones et al., 2018) and mushroom body markers from single-cell sequencing in *Drosophila* (Davie et al., 2018) and *Apis mellifera* (Traniello et al., 2020) were taken from supplemental information of published work; for *Apis mellifera*, mushroom body clusters were defined as the clusters with *mub* as a marker gene.

### *Drosophila* single-cell sequencing

For **Fig. 5H**, Single-cell sequencing from *Drosophila* (Davie et al., 2018) was downloaded from the scope.aertslab.org website. The x and y positions of each cell on the tSNE were specified by this object, and the position of ensheathing glia indicated was informed by the cluster identities defined in Davie et al. The expression level of *wdp* was computed by normalizing the expression levels provided; taking the log2 of UMIs normalized for the total UMI in each cell and multiplied by a scaling factor of 10,000.

### Synteny analysis of *wdp* locus

Corresponding genes between *Drosophila, Apis mellifera*, and *Harpegnathos* were found using a BLASTp search; each *Harpegnathos* gene, excluding the lncRNA LOC112590028, had a best match to the gene indicated in the other two species.

## Data Availability

Next generation sequencing data generated for this study have been deposited in the NCBI GEO with accession number GSE172309. Sequencing data will remain private during peer review and released upon publication, but will be made available upon request.

## Acknowledgments

The authors thank Katy Munson and Alexandra Mackenzie at the UW PacBio Sequencing service for performing Iso-Seq; Jakub Mlejnek for technical support; Janko Gospocic for helpful discussion; Claude Desplan and Danny Reinberg for support and encouragement. R.B. was supported in part by the NIH (DP2MH107055, R01AG071818). LD and BS were supported by NIH grant R01 AG058762 from the National Institute on Aging; B.S. was supported by the Long-Term Fellowship LT000010/2020-L from the Human Frontier Science Program.

---

**Figure S1.**
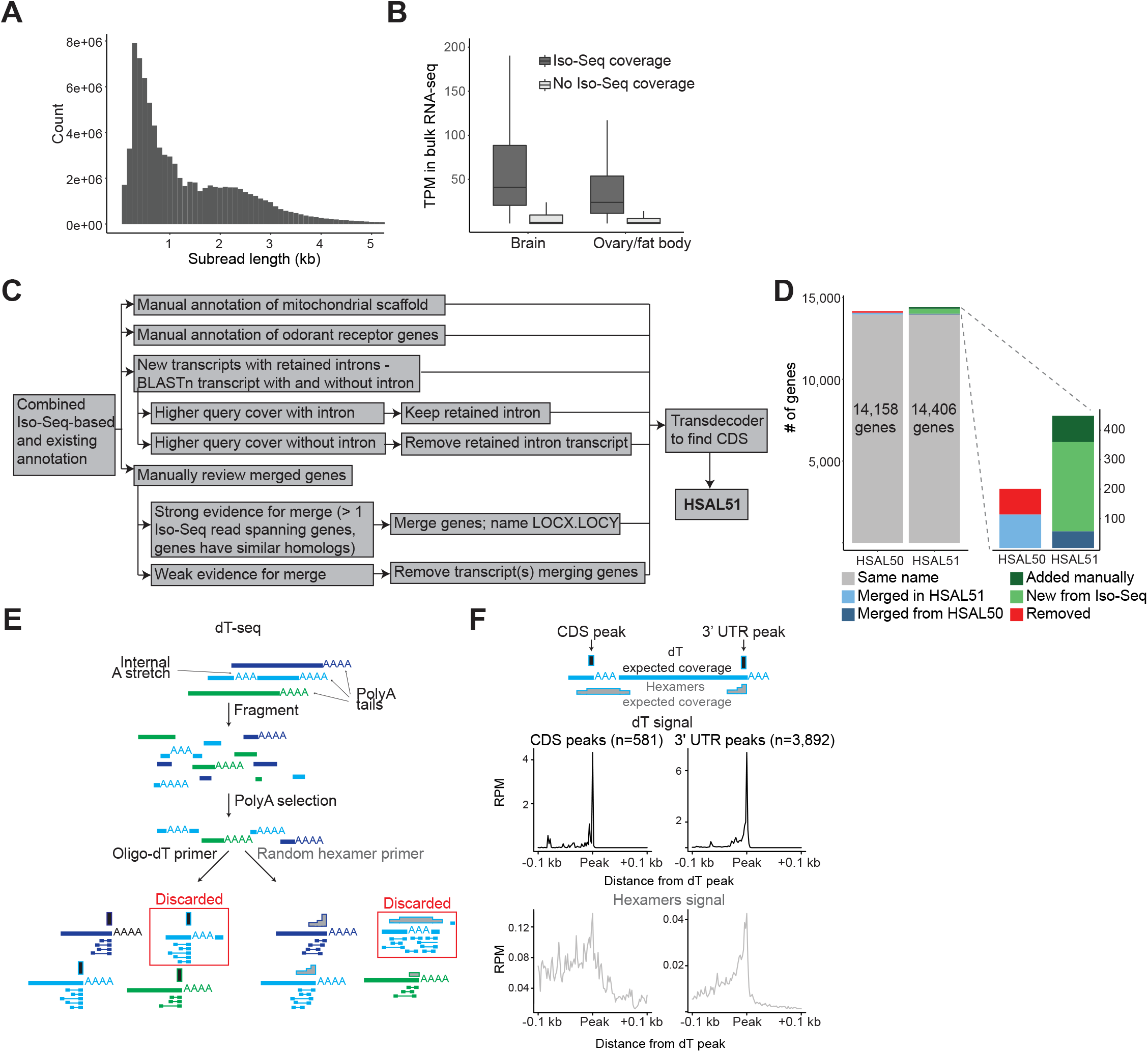
Statistics and methods used to create and evaluate the new *Harpegnathos* annotation. (A) Length distribution of all raw Iso-Seq subreads. (B) Transcripts per million (TPM) from short-read RNA-seq of genes with and without Iso-Seq coverage in brain and fat body/ovary. (C) Pipeline for manual annotation following combination of Iso-Seq-based and RNA-seq-based annotations. (D) Relationship between gene models in HSAL50 and HSAL51. (E) Schematic of the dT-seq approach. RNA was chemically fragmented. PolyA+ molecules were purified and split into two reverse transcription reactions, one primed with an anchored oligo-dT primer and one with random hexamers. The resulting cDNA was assembled into libraries and sequenced. The scheme at the bottom shows that the expected read distribution in dT-and hexamer-primed reactions differs for true polyA tails and internal A-stretches. This information was used to discard peaks that did not correspond to *bona fide* TTS (red square). (F) Expected (top) and observed (bottom) signal at dT peaks found in the CDS (let) or 3’ UTR (right) from oligo-dT primed libraries (”dT”, top) and random hexamer primed libraries (”hexamers”, bottom).

**Figure S2.**
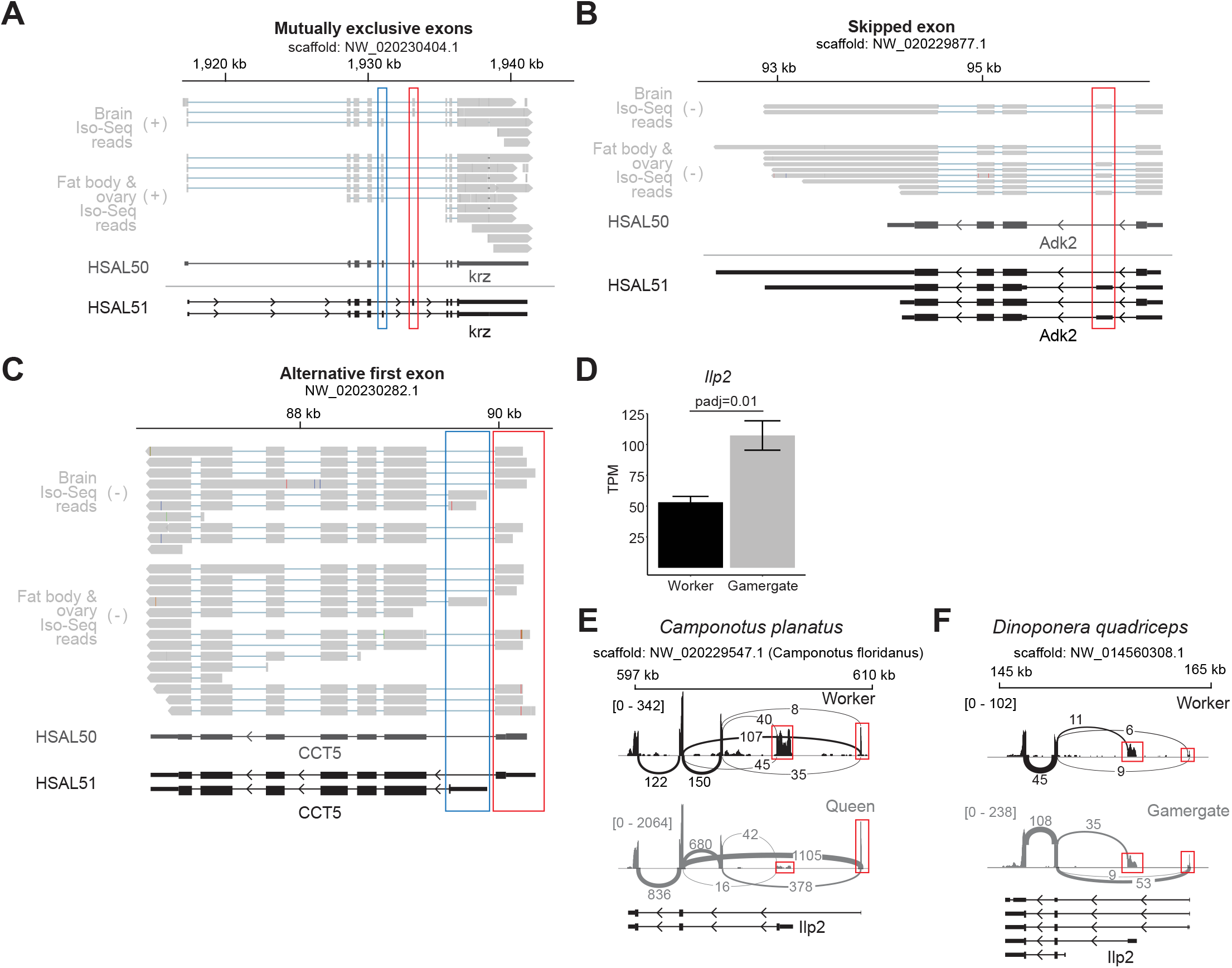
More comparisons of alternative splicing in HSAL50 and HSAL51. (A-C) Examples of a transcript with newly identified alternative splicing patterns of (A) mutually exclusive exons, (B) a skipped exon, and (C) an alternative first exon. Boxes indicate region of the gene that is alternatively spliced. A subset of HSAL51 isoforms is shown. (D) TPM of *Ilp2* (*LOC105188195*) in worker (n=11) and gamergate (n=12) brains. P-adjusted is from DESeq2 differential expression analysis. (E) Sashimi plot for the *Ilp2* gene (*LOC105257206*) in *Camponotus planatus* (RNA-seq from Chandra et al., 2018; using *Camponotus floridanus* genome and annotation) for worker (n=5) and queen (n=5) brains. Splice junction line widths are scaled to the number of reads spanning the splice junction and the total number of reads mapped to *Ilp2* for each caste. Red boxes indicate positions of first exon for each isoform. (F) Sashimi plot for the *Ilp2* gene (*LOC106750697*) in *Dinoponera quadriceps* (RNA-seq from Patalano et al, 2015) for worker (n=6) and gamergate (n=6) brains. Splice junction line widths are scaled to the number of reads spanning the splice junction and the total number of reads mapped to *Ilp2* for each caste. Red boxes indicate positions of first exon for the two major isoforms.

**Figure S3.**
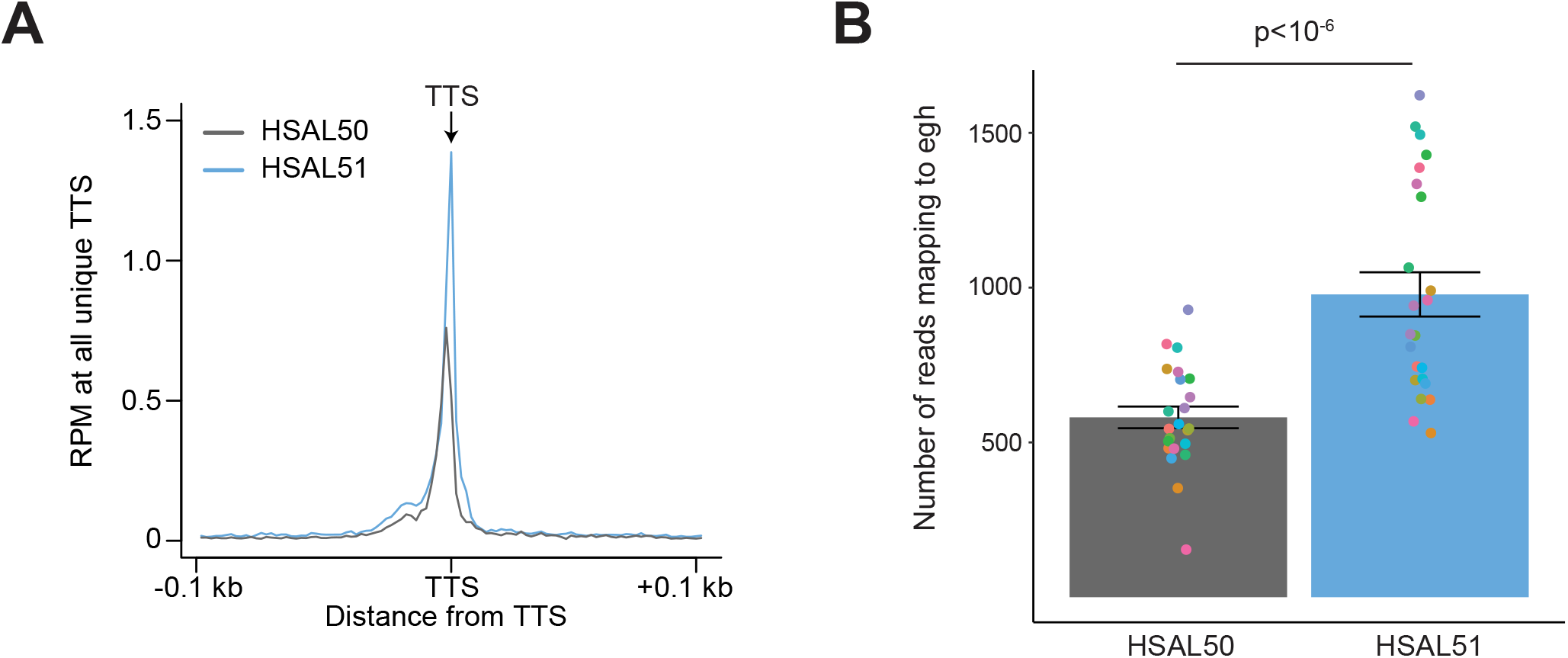
Transcript extensions and RNA-seq analysis. (A) dT-seq coverage (see Fig. S1E and methods) coverage at all unique TTS in HSAL50 (gray) and HSAL51 (blue). (B) Reads mapping to *egh* in HSAL50 and HSAL51. P-value is from a paired Student’s t-test.

**Figure S4.**
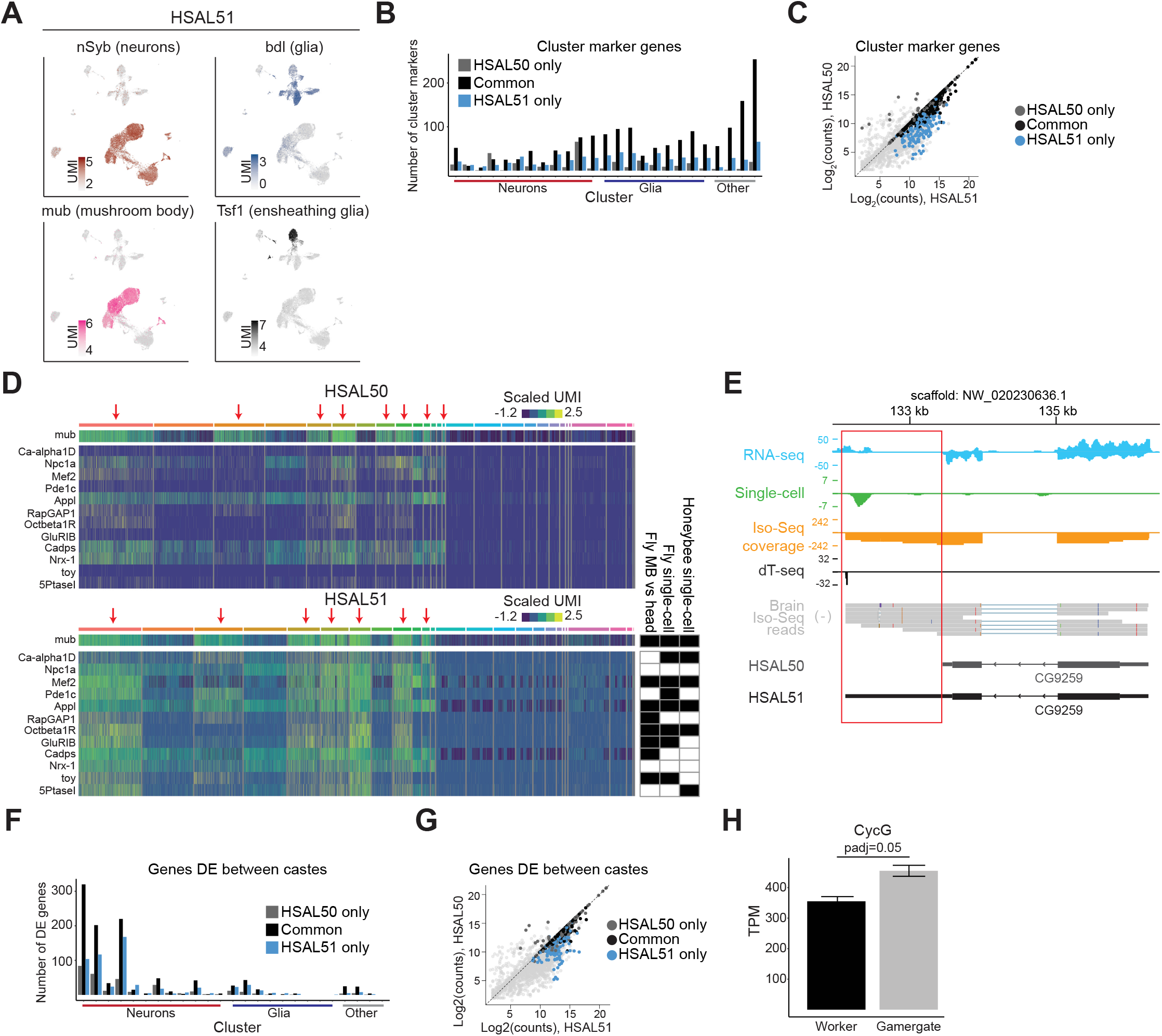
Additional single-cell analyses using HSAL50 and HSAL51 annotations. (A) Heatmaps of markers for neurons (*nSyb*), glia (*bdl*), mushroom body neurons (*mub*), and ensheathing glia (*Tsf1*) in HSAL51 single-cell clustering. (B) Number of marker genes (p-adjusted < 0.05, LFC > 1) for each cluster. Marker genes common to HSAL50 and HSAL51 analyses are shown in black, while markers unique to HSAL50 are in gray and markers unique to HSAL51 are in blue. (C) Scatter plot for UMI counts in HSAL50 vs. HSAL51 with marker genes highlighted according to (B). (D) Heatmap of newly identified mushroom body markers in HSAL51 (padj < 0.05 and logFC > 1.5). Arrows denote mushroom body clusters, as determined by *mub* expression (top row of heatmap). Classification of each new marker in other data sets (fly MB vs head, (Jones et al., 2018); fly single-cell, (Davie et al., 2018); honeybee single-cell, (Traniello et al., 2020)) is indicated in heatmap to right, with black boxes indicating marker was identified as mushroom body-enriched (see methods). (E) Genome browser view showing *CG9259* with RNA-seq, Iso-Seq, and single-cell coverage along with dT-seq and raw Iso-Seq reads. Scales represents counts per million. A subset of HSAL51 isoforms is shown. (F) Number of genes differentially expressed (DE) within each cluster (padj < 0.01). Differentially expressed genes common to HSAL50 and HSAL51 analyses are shown in black, while genes unique to HSAL50 are in gray and genes unique to HSAL51 are in blue. (G) Scatter plot for UMI counts in HSAL50 vs. HSAL51 with differentially expressed genes highlighted according to (F). (H) TPM of *CycG* in worker (n = 11) and gamergate (n = 12) brains. P-adjusted is from DESeq2.

**Figure S5.**
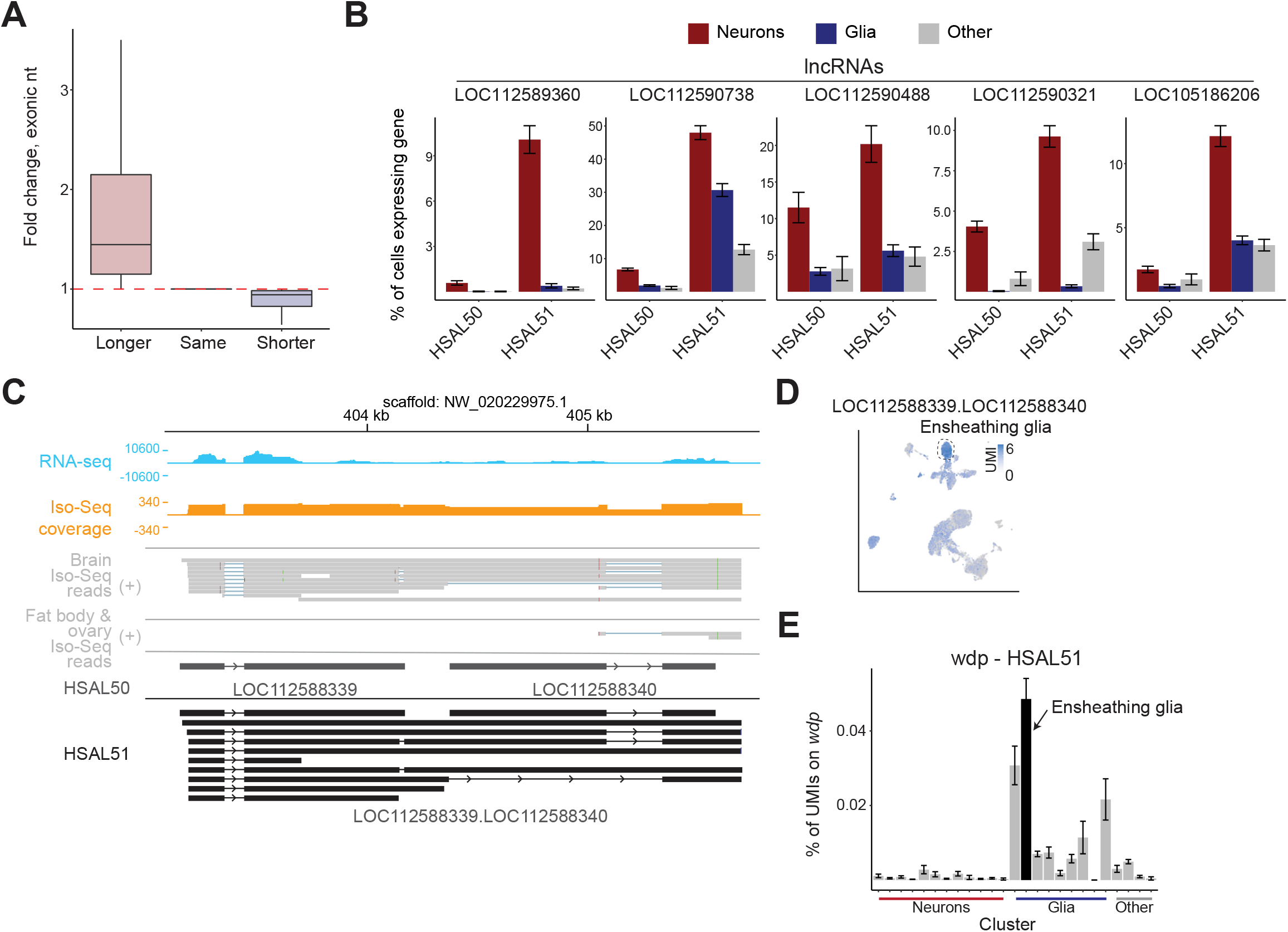
Additional single-cell analyses of lncRNA expression. (A) Fold-change in nucleotides covered by exons of lncRNAs for each category in Fig. 5A. (B) Examples of neuronal lncRNA detected in HSAL51 showing % of neurons, glia, and other cells expressing the indicated genes in HSAL50 and HSAL51. (C) Genome browser view showing RNA-seq signal, Iso-Seq signal, and raw Iso-Seq reads of a the locus containing two ensheathing glia marking lncRNAs which were merged into one gene model in HSAL51. Scales for RNA-seq and Iso-Seq represent counts per million. (D) Heatmap showing normalized UMI counts of the new merged lncRNA *LOC112588339*.*LOC112588340* in HSAL51. (E) % of UMIs mapping to *wdp* in HSAL51.

